# Tumor Microenvironment Alters Chemoresistance of Hepatocellular Carcinoma Through CYP3A4 Metabolic Activity

**DOI:** 10.1101/2021.01.23.427874

**Authors:** Alican Özkan, Danielle L. Stolley, Erik N. K. Cressman, Matthew McMillin, Sharon DeMorrow, Thomas E. Yankeelov, Marissa Nichole Rylander

**Author notes:** **Correspondence:** Alican Özkan.

## Abstract

Variations in tumor biology from patient to patient combined with the low overall survival rate of hepatocellular carcinoma (HCC) present significant clinical challenges. During the progression of chronic liver diseases from inflammation to the development of HCC, microenvironmental properties, including tissue stiffness and oxygen concentration, change over time. This can potentially impact drug metabolism and subsequent therapy response to commonly utilized therapeutics, such as doxorubicin, multi-kinase inhibitors (e.g., sorafenib), and other drugs, including immunotherapies. In this study, we utilized four common HCC cell lines embedded in 3D collagen type-I gels of varying stiffnesses to mimic normal and cirrhotic livers with environmental oxygen regulation to quantify the impact of these microenvironmental factors on HCC chemoresistance. In general, we found that HCC cells with higher baseline levels of cytochrome p450-3A4 (CYP3A4) enzyme expression, HepG2 and C3Asub28, exhibited a cirrhosis-dependent increase in doxorubicin chemoresistance. Under the same conditions, HCC cell lines with lower CYP3A4 expression, HuH-7 and Hep3B2, showed a decrease in doxorubicin chemoresistance in response to an increase in microenvironmental stiffness. This differential therapeutic response was correlated with the regulation of CYP3A4 expression levels under the influence of stiffness and oxygen variation. In all tested HCC cell lines, the addition of sorafenib lowered the required doxorubicin dose to induce significant levels of cell death, demonstrating its potential to help reduce systemic doxorubicin toxicity when used in combination. These results suggest that patient-specific tumor microenvironmental factors, including tissue stiffness, hypoxia, and CYP3A4 activity levels, may need to be considered for more effective use of chemotherapeutics in HCC patients.

## 1 Introduction

Cancer is the second-highest cause of mortality in the United States, lagging just slightly behind cardiovascular disease in 2020 ^1^. Among all cancer types, hepatocellular carcinoma (HCC) has the second-lowest 5-year survival rate (17.7%) and has shown the highest increase in mortality among all cancers over the past seven years ^2^,^3^. Complicating treatment, HCC is commonly diagnosed at an intermediate or an advanced stage and often occurs secondary to underlying chronic liver disease and cirrhosis ^4^. The prognosis is poor as treatment options are limited by compromised liver function due to underlying disease. Despite screening efforts for at-risk patients, most are not surgical candidates for partial resection, and the availability of full liver transplantation is very low relative to the need^5^.

These issues mean that systemic and localized drug-based therapies play a significant role in current standard therapy for HCC. Despite these therapeutic interventions, the survival rate for HCC remains low, partially attributed to the variable efficacy of current treatment methods based on underlying factors ^6–8^. As such, stratifying patients for the most effective treatment is critical because of three factors; the degree of tumor burden, the degree of liver dysfunction, and highly variable treatment efficacy between patients ^9^. For example, the tyrosine kinase inhibitor sorafenib has shown modest success in selected patients as a systemic treatment ^10^, but effectiveness is tempered by poor tolerance of the drug in many instances^9^. Localized delivery of drugs through transarterial chemoembolization (TACE) combines delivery of drugs such as doxorubicin with embolization to promote localized ischemia and hypoxia. TACE blocks the arterial blood supply of a tumor through particulate or viscous liquid agents such as degradable starch microspheres, drug-eluting beads, or ethiodized oil. This is a well-established technique that allows a high local dose while simultaneously increasing the residence of chemotherapeutic drugs in the target area, cutting off the supply of nutrients, and also limiting exposure and toxicity for the rest of the body ^11^. This has emerged as the standard of care for intermediate-stage HCC. However, tumor cells in the hypoxic environment may undergo phenotypic adaptations that aid survival. Such changes may account for the high rate of persistent, viable tumor cells observed after TACE in previous studies ^12^. Therefore, while a substantial survival benefit can be realized, there is still much room for improvement and understanding of the changes that occur in the tumor cells during embolization^6–8^.

It is well established that many of the difficulties in treating HCC may stem from the numerous tumor microenvironment (TME) changes that occur in underlying chronic liver disease and the rapid progression of HCC. The modulation of the TME has been shown to impact drug metabolism significantly and is thought to be a major contributor to the known differential response of patients to chemotherapy ^13,14^. Furthermore, induction of hypoxia in the TME due to stiffening of the extracellular matrix (ECM) and embolization during treatment can alter the chemoresistance of the tumor cells, further impacting the treatment efficiency in intermediate and advanced stage HCC ^15–17^. The modulation of response and the individual impact of these TME features have yet to be fully characterized in a three-dimensional (3D) HCC-TME model.

The increase in microenvironmental stiffness resulting from fibrosis, usually culminating in cirrhosis, is a hallmark of most chronic liver diseases and is observed in 80-90% of HCC patients ^4^. The most notable hallmark of liver cirrhosis that impacts cellular and tissue function is increased collagen deposition from activated hepatic stellate cells, increasing the stiffness and compression modulus of liver tissue ^18^. Similarly, the progression of HCC is marked by further localized stiffening. This desmoplastic reaction is attributed to further differentiation of liver stellate cells into myofibroblasts ^18^, resulting in additional deposition of collagen^19^. This stiffening of the ECM, in conjunction with high tumor cell density, further reduces local oxygen and nutrient diffusion. Limited oxygen availability has been shown to alter the outcomes of the chemotherapeutic treatments by affecting the drug transporters’ p-glycoprotein (MDR1, multidrug resistance 1), drug targets (topoisomerase II), or by initiating drug-induced apoptosis ^20^. Subsequent alterations in the cancer cells’ response to chemotherapy can occur through modulation of chemoresistance markers and hepatocyte metabolic enzymes such as cytochrome P450 (CYP450), primarily the CYP3A4 subgroup ^21^. Thus CYP3A4 expression can potentially serve as an indicator for predicting chemotherapeutic response ^22^. This highlights a potential mechanism of differential tumor chemoresistance through the modulation of CYP3A4 under different microenvironmental conditions.

Numerous *in vitro* models have been used to study the impact of TME modulation on HCC treatment response. Two-dimensional (2D) cell monolayers have documented an increase in HepG2 cell survival following exposure to doxorubicin during hypoxia compared to normoxic conditions ^23^. However, traditional 2D cell culture models do not allow for adequate representation of the physiological diffusion and associated transport barriers found in the 3D extracellular microenvironment, limiting clinical translation. Tumor spheroid models have been utilized as a more representative system for assessing HCC drug response in direct cell-cell contacts and the subsequent decrease of HCC chemoresistance ^24^. However, the lack of ECM in these models severely curtails the study of the interactions with and subsequent tuning of the ECM components of the TME ^25^.

Previous efforts that address the importance of ECM microenvironmental properties on the chemotherapeutic response, employ tunable hydrogels models, composed of cells cultured in ECMs of collagen, fibrin, alginate, and Matrigel™. These models have been used to investigate the ECM’s role in *in vitro* studies of chemotherapy response, drug transport, cell invasion, and differentiation ^25,26^. Culturing different breast cancer cell lines in an alginate hydrogel showed doxorubicin chemoresistance was altered in particular phenotypes when ECM stiffness was increased ^27^. With respect to metabolic activity, one study investigated Ifosfamide metabolism by C3A HCC cells with different levels of CYP3A4 expression when cultured in polylactic acid (PLA) ^28^. However, this study reported treatment efficacy only with glioblastoma cells and did not explore or discuss the treatment response of liver cells. Another study demonstrated that culturing Caco-2 colorectal cancer cells on the top of a 3D collagen-Matrigel™ blended hydrogel with dynamic flow conditions increased CYP3A4 expression drastically compared to 2D monolayers without flow ^29^. Similarly, another study showed that the CYP3A4 activity of HepaRG HCC cells increased when cells were cultured in hyaluronan-gelatin or wood-derived nanofibrillar cellulose ECMs relative to 2D monolayer culture ^30^. Research has shown culturing U251 and U87 glioblastoma cells in 3D PLA scaffold under hypoxia exhibited higher resistance to doxorubicin and greater production of basic fibroblast growth factor (bFGF) and vascular endothelial growth factor (VEGF)^31^. Furthermore, U87, U251, and SNB19 glioblastoma cells have been shown to be more resistant to temozolomide when cultured in scaffold-free spheroids under hypoxic conditions compared to comparable spheroids under normoxic conditions ^32^. Other studies have used native ECM (collagen and fibrin) and non-native polymers (agar, acrylamide, and polylactic acid) to recapitulate 3D breast and hepatic tumor microenvironments. However, these studies did not investigate the regulation of cellular metabolism, including CYP3A4 activity, under different microenvironmental conditions ^26,27,33,34^. Recent work extending these efforts has demonstrated significant potential in utilizing collagen tunability to replicate native microenvironment properties (pH, stiffness, fiber properties, and porosity) ^35,36^, demonstrating promise in replicating physiological TME to investigate the modulation of CYP3A4 activity and chemoresistance.

In this study, we utilized a collagen type I hydrogel to investigate the influence of ECM stiffness, oxygen concentration, cell type, and the availability of a 3D microenvironment (2D vs. 3D) on drug metabolism and response to two common chemotherapeutic agents, doxorubicin and sorafenib, both individually and in combination. We used four established HCC cell lines (HepG2, C3Asub28, HuH-7, and Hep3B2), which present with different basal metabolic profiles of CYP3A4 expression to quantify the impact of modulations of the TME. Our results demonstrate the importance of the contribution of a 3D ECM in drug design and dosing based on the significant differences seen in drug response and metabolism when tumor cells are cultured in 3D collagen type I compared to traditional 2D culture. Further, we show that variations in the TME, including liver stiffness and hypoxia, results in altered drug metabolism and subsequent drug efficacy. Tissue stiffness, varied by using collagen concentrations comparable to normal and cirrhotic liver stiffnesses^37^, caused an alteration in chemoresistance and drug metabolism. TME oxygen regulation to simulate normoxic and hypoxic conditions produces a similarly significant, general effect on both chemoresistance and drug metabolism. However, the TME regulation was not consistent for every cell type investigated, shown in the heterogeneous regulation of cell chemoresistance and drug metabolism in the HCC cell lines. This highlights a potential clinically translational impact of HCC genetic polymorphisms and different etiologies on treatment outcome. Specifically, we identify that the basal cellular CYP3A4 metabolism can be differentially regulated by TME hypoxia and tissue stiffness, thus impacting the efficacy of commonly used HCC therapeutics. This relationship provides a potential explanation of the poor outcomes of drugs in HCC clinical trials and may eventually lead to improve outcomes for HCC patients.

## 2 Methods

### 2.1 Cell Culture

Human hepatocellular carcinoma cell lines HepG2 (HB-8065™, ATCC^®^, Manassas, VA), HuH-7, Hep3B2.1.7, and HepG2 derived C3A with enhanced expression of CYP3A4 mRNA and CYP3A4-mediated activity (C3Asub28) ^38^ were used in this study. HuH-7 cells express a mutated form of tumor-suppressive protein p53, leading to an increased half-life and accumulation of the protein in cell nuclei. This has been shown to correlate with increased chemoresistance ^39^. Hep3B2.1.7 was used as an example of an HCC tumor with hepatitis B DNA in the genome and subsequent mutations, including partially deleted and suppressed expression of p53. All cells were cultured with DMEM supplemented with 10% heat-inactivated fetal bovine serum (FBS, F4135, Sigma Aldrich, MO) and 1% Penicillin/Streptomycin (P/S, Invitrogen, CA). Normoxic conditions were similarly generated by culturing cells in standard cell culture conditions in a normoxic incubator (21% O_2_, 5% CO_2_, 37°C Thermo Fisher Scientific, Rochester, NY, USA). Hypoxic conditions were simulated by placing cells in a sealed incubator (1% O_2_, 5% CO_2_). All cells were grown to approximately 70% confluence and used within the first eight passages.

### 2.2 Preparation and Tuning of Collagen

Type I collagen isolated from rat tail tendons (donated by the University of Texas at Austin - Institute for Cellular and Molecular Biology) was used to recapitulate the tissue microenvironment as it is the primary ECM component of human tissue, including the liver ^26^. As we have previously published, a stock solution of type I collagen was prepared by dissolving excised rat tail tendons in an HCl solution at a pH of 2.0 for 12 hours at 23°C ^26^. The solution was then centrifuged at 4°C for 45 minutes at 30000 g, and the supernatant was collected, lyophilized, and stored at −20°C. The lyophilized collagen was mixed with diluted 0.1% glacial acetic acid, maintained at 4°C, and vortexed every 24 hours for three days to create a collagen stock solution. Finally, collagen was centrifuged at 4°C for 10 minutes at 2700 rpm to remove air bubbles. Collagen concentrations of 4 and 7 mg/ml were used to replicate normal and cirrhotic liver stiffness, respectively, which we have previously demonstrated to match native liver compression moduli after cells have reached their native morphology ^26,37^. Collagen solutions were adjusted to pH 7.4 with 1X DMEM, 10X DMEM (Sigma Aldrich, St. Louis, MO), and 1N NaOH (Fisher Scientific, Pittsburgh, PA.). Following this, the collagen mixture was mixed with the intended HCC cell lines uniformly at a concentration of 1×10^6^ cells/ml. Each suspension was dispensed as 50 μL aliquots in 96 well plates and allowed to polymerize. For 2D monolayer samples, identical numbers of cells in 100 μL of media were dispensed into the wells of a 96 well plate. Cell media was changed every two days.

### 2.3 Confined Compression Test

Cirrhotic stiffening in the TME has been shown to alter the chemoresistance of many cancer cell types ^40^. As a result, the mechanical properties of the TME, such as compression modulus, also increase. The compression modulus of 3D collagen hydrogels was measured with quasi-steady uniaxial unconfined compression (Instron, Norwood, MA) ^26^ to ensure there is no significant difference between collagen gels with different HCC cell lines three days after seeding. In the analysis, the hydrogels are assumed to be linear under the deformation conditions, and the slope of the stress-strain curve represents the compression modulus. Hydrogels were prepared as described in the previous section, and 500 μL of hydrogels were placed inside 24 well plates. Polymerized collagen samples were punched (9.53 mm diameter) to remove the concave meniscus at the sample edge. Samples were compressed using a 20 mm diameter load cell of a flat steel surface. Load cells (10 N Static Load Cell, 2519-10N) were lowered approximately 2.5 mm away from the flat surface and displaced 2 mm at a rate of 0.0085 mm/s to achieve 0.1% strain/s over the range of 0-20% strain. Stress was calculated from the force response divided by the initial area of collagen sample (71.26 mm^2^). All measurements were performed at room temperature (23°C) and the total duration of each experiment was less than 4 minutes. The data was analyzed using Matlab^®^ (MathWorks, Natick, MA).

### 2.4 Dosing With Chemotherapeutics

Doxorubicin and sorafenib have been commonly used in ongoing clinical trials for HCC treatment either alone or in combination ^41^. Prepared samples were exposed to doxorubicin (D1515, Sigma-Aldrich, St. Louis, MO) with or without sorafenib (HY-10201, MedChemExpress LLC, Monmouth Junction, NJ,) for 24 and 48 h. A broad range (1 nM – 200μM) of doxorubicin concentrations were used to determine the response across the different HCC cell lines. Three different doses of sorafenib were used to replicate previously tested effective concentrations: none (0 μM), standard (11 μM), and high (22 μM) ^42^. Samples were washed with warm (37°C) 1x phosphate buffered saline (PBS) three times to remove the excess drug after the treatment. Cell culture media was used as a negative control.

#### Measuring Viability

Cell viability 72 hours after the completion of drug treatment was measured with Cell Titer Blue (G8081, Promega, Fitchburg, WI) cell viability assay to quantify the response of HCC cells to chemotherapeutic treatment under varying TME conditions (such as stiffness and hypoxia). Briefly, cell media was mixed at a ratio of 5:1 with assay solution and incubated at 37°C for 1 hr. A Cytation 3 plate reader was used to read fluorescence (Ex: 530 nm/Em: 620 nm) of the assay. Findings were normalized to control (cells treated with drug-free media) to obtain percent cell viability. The data gathered from cell viability assays after doxorubicin treatment was fitted using the cftool function of Matlab^®^ (MathWorks, Natick, MA) and half-maximal inhibitory concentration (IC_50_) value was calculated and reported as chemoresistance indicator. As sorafenib has been used as a prodrug used for angiogenesis treatment, mainly by inhibiting vascular endothelial growth factor (VEGFR), platelet-derived growth factor receptor (PDGFR), and rapidly Accelerated Fibrosarcoma (RAF), but it also has a minor direct cytotoxic side-effect in some instances ^15^. For that reason, treatment efficacy of standalone sorafenib and combined treatments were reported as fold viability change compared to untreated control for the two clinically relevant doses used, standard (11 μM), and high (22 μM). Additionally, for combined treatment, the minimum required doxorubicin dose to initiate toxicity was reported. For this analysis, statistical comparison (p<0.05) of viability under the tested doses and untreated samples were compared, and the minimum dose was reported.

### 2.5 Modeling of Oxygen Consumption in Comsol Multiphysics

To verify 3D collagen gels do not induce hypoxia in the system, we modeled oxygen concentration across the culture media and collagen gel. Depletion of oxygen in collagen hydrogels by HCC cells were modeled using Comsol Multiphysics (Comsol Inc), which is a finite element analysis solver software. For this analysis, the computational domain was assumed to be 2D and axisymmetric at the center of the hydrogel and culture medium (Figure 1a). Modeling parameters are defined and parameter values are provided in Table S-I. 25000 domain elements were added to solve the problem in the computational domain. Accordingly, time dependent convective diffusion equations in transport of diluted specifies module were solved temporally over the collagen hydrogel (Equation 1) and culture medium (Equation 2):

**Figure 1:**
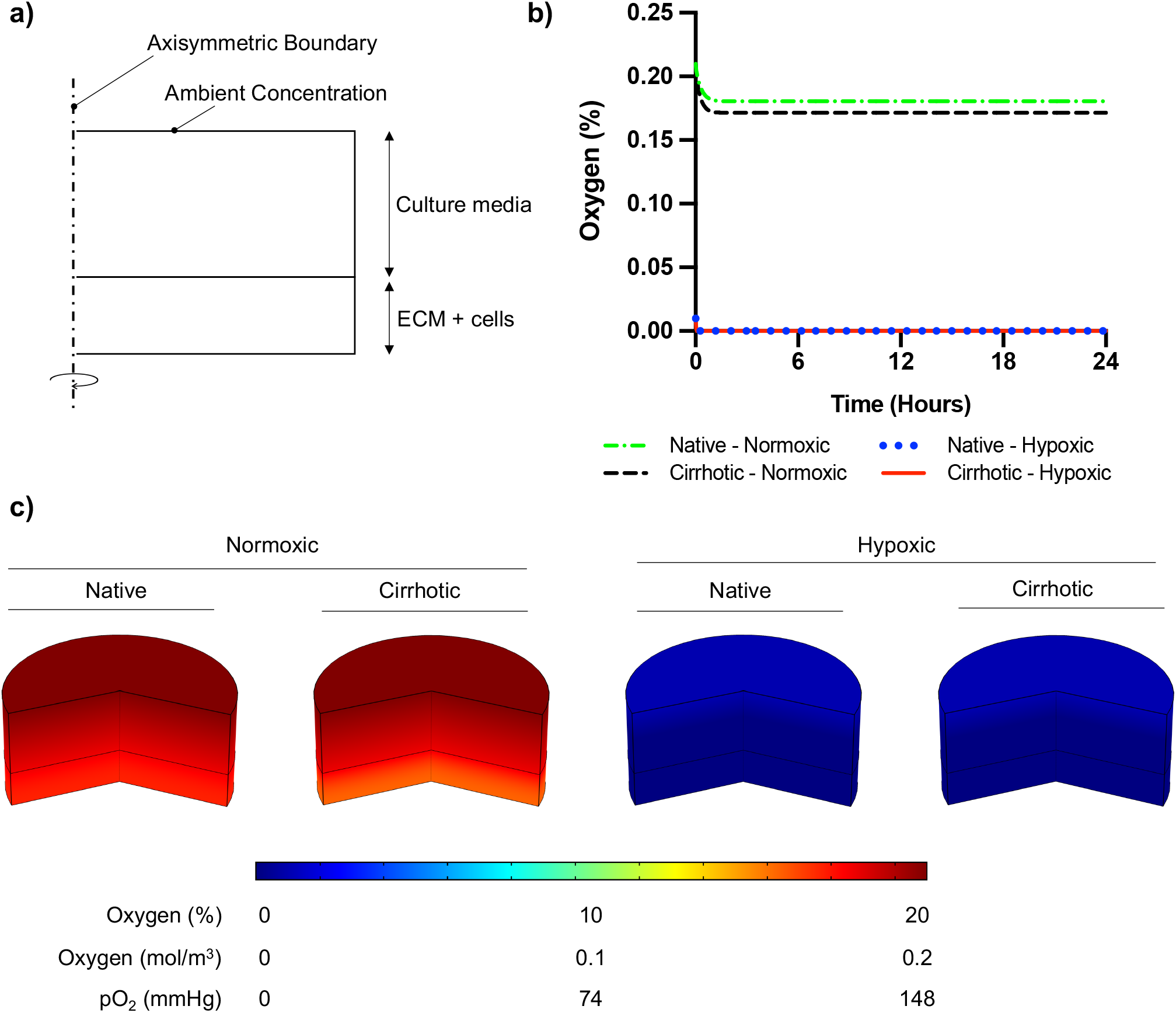
Oxygen depletion modeling in ECM and culture media. a) Computational domain of the problem in 2D axisymmetric configuration. b) Temporal average oxygen concentration in collagen hydrogels. Oxygen concentration distribution reaches steady state less than one hour after cell seeding. No marginal variation was observed in oxygen concentrations between native and cirrhotic hydrogels. c) 3D contour of oxygen concentration across the collagen hydrogel and culture media after system reaches steady state.

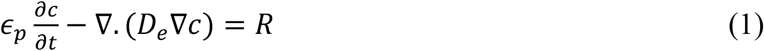

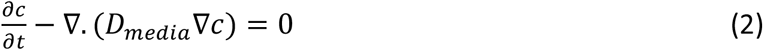

where c is the oxygen concentration, ϵ_*p*_ is porosity of the collagen hydrogel, *ρ* is the density of media, heat capacity of media, *D*_*e*_ is effective oxygen diffusivity, R is oxygen consumption rate by HCC cells. The diffusivity of collagen hydrogels was adjusted using Bruggemen model, where the porosity of collagen hydrogels was required to be implemented. Accordingly, Equation 3 was incorporated to the computational model:

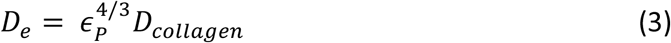

### 2.6 CYP3A4 Activity Measurement

Overexpression of CYP3A4 has been shown to decrease the efficacy of chemotherapeutics and is one of the main challenges in the treatment of HCC tumors ^22^. To relate the chemotherapeutic response to the metabolic activity of HCC cells under the influence of hypoxia and different stiffnesses, the expression of CYP3A4 was quantified using P450-Glo™ Assay and Screening Systems (V9001, Promega, Madison, WI) in response to different oxygen concentrations, normoxic and hypoxic, and different spatial conditions, 2D, 3D normal, and 3D cirrhotic. HCC cells were cultured in 2D or 3D, as previously described for three days before CYP3A4 measurements to allow cells to reach native morphology ^37^. Prepared samples were washed twice with PBS and incubated with 50 μL of CYP3A4 substrate Luciferin-IPA (3 μM, dissolved in 1X DMEM). After 1 hour of incubation at 37°C, 50 μL of Luciferin Detection Reagent was added and pipetted up and down several times to ensure cell lysis. After 20 minutes of incubation at room temperature, cell supernatants were transferred to a 96-well opaque white luminometer plate (white polystyrene; Costar, Corning Incorporated) and luminescence was measured using a Cytation 3 plate reader (BioTek Instruments, Inc., VT). Reagents without cells were included as background controls. Metabolic activities were calculated by subtracting the background luminescence and normalizing to the seeded cell number.

### 2.7 Statistical Analysis

Two-tailed student’s t-test assuming unequal variance was performed in Matlab^®^ (MathWorks, Natick, MA) to compare samples, and a p-value less than 0.05 was considered significant for variation. Pearson correlation between CYP3A4 expression and IC_50_ findings was performed in Graphpad Prisim (Graphpad Holdings, LLC). Data are reported as mean ± standard deviation unless otherwise indicated. All experiments were replicated a minimum of four times.

## 3 Results

HCC cells were cultured for three days to reach native morphology in a monolayer in a tissue culture plate (2D) or in rat tail-derived collagen type I hydrogels (3D). Cells were treated with doxorubicin with or without sorafenib for 24 or 48 hours. Viability was assessed 72 hours after the end of drug treatment, as described in Figure 2b. Before doing so, we simulated oxygen consumption in collagen hydrogels to observe weather cirrhosis alters oxygen concentration in hydrogels as presented in Figure 2.

**Figure 2:**
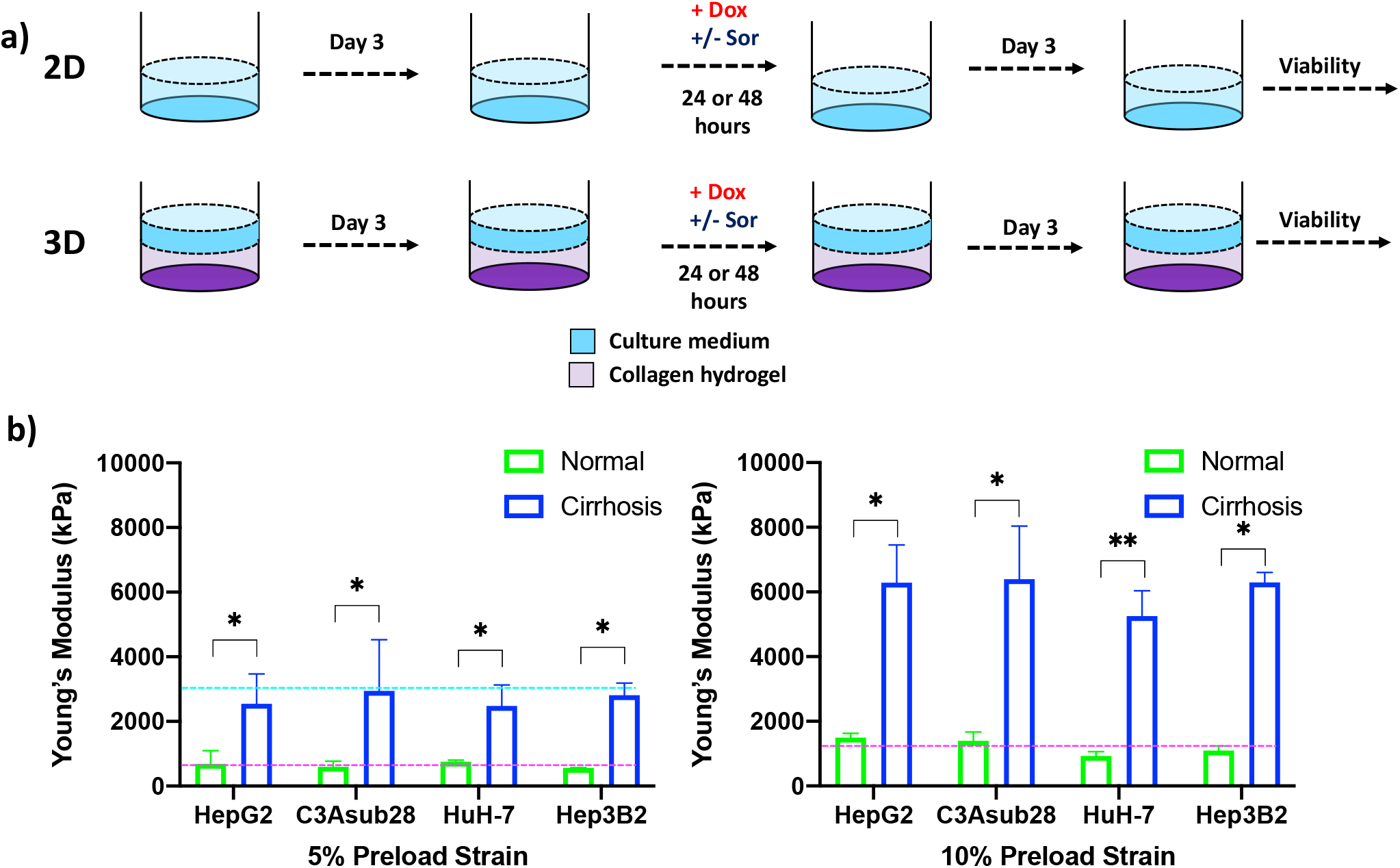
HCC cells cultured in a monolayer or in collagen hydrogels of varying stiffness. **a)** Experimental procedure outline. Cells were allowed to adhere and reach their native morphology before the treatment for 24 or 48 hours. **b)** Compression modulus of the HCC hydrogels with 4 and 7 mg/ml collagen concentrations, which replicate normal and cirrhotic tissues, respectively. The compression modulus did not vary when different HCC cells were cultured in collagen hydrogels. Compression modulus values were significantly different when different collagen concentrations were used to replicate normal and cirrhotic conditions. Dashed lines represent patient compression modulus of healthy (magenta) and cirrhotic (magenta) tissues. ^43,44^. Selected collagen concentrations replicated native tissues successfully at different preload strains. * p<0.05, ** p<0.01.

### 3.1 Oxygen concentration in Collagen Hydrogels

To confirm 3D collagen gels do not introduce hypoxia in the system, oxygen concentration was modeled across the gel and culture media. According to Comsol Multiphysics simulation results presented in Figure 1b, oxygen concentration slightly decreases in collagen hydrogels under normoxic culture conditions. Under normoxic culture conditions, oxygen concentration in cirrhotic collagen gel decreased to 17.15%. Oxygen concentration in normoxic culture conditions with native collagen gel resulted in 18.06%, which is slightly higher than cirrhotic conditions. The drop in oxygen concentration is a result of standardized polystyrene well plates not being gas permeable. Nevertheless, the oxygen drop compared to the initial concentration is minimal and we observe a marginal variation between native and cirrhotic hydrogels in normoxic conditions. Under hypoxic conditions, we observe depleted and uniform oxygen in hydrogels. Accordingly, we can state that oxygen concentration was uniform across the hydrogels and oxygen difference between native and cirrhotic conditions was not different (Figure 1c).

### 3.2 Native 3D Microenvironment Compression Modulus

Hepatocellular carcinoma cells uniformly embedded inside rat tail-derived collagen type I hydrogels were allowed to reach their native morphology for three days. Afterwards, stiffness of the collagen hydrogels was quantified using a uniaxial compression test with 5% and 10% preload strains as presented in Figure 2b to determine any potential impact of different HCC cell lines on collagen stiffness. Our results demonstrate that increasing collagen concentration significantly elevated the compression modulus of the collagen gels (p<0.05). However, we found no significant difference between compression modulus between collagen hydrogels with the different HCC cell lines over the timeframe considered. Collagen hydrogels at 4 mg/ml concentration produce a compression modulus comparable to tissue in a normal hepatic microenvironment, which has been reported to be at 0.64 ± 0.08 kPa and 1.08 ± 0.16 kPa for 5% and 10% preload strains, respectively ^43^. At 4 mg/ml collagen concentration, the average compression modulus was found to be 0.66 ± 0.07 kPa and 0.11 ± 0.01 kPa for 5% and 10% preload strains, respectively. Likewise, at 7 mg/ml collagen concentration, collagen hydrogels achieved a compression modulus comparable to a human hepatic tumor microenvironment, which has been reported to be 3 kPa under 5% preload strain ^44^. The significant difference (p<0.05) of compression moduli between 7 mg/ml and 4 mg/ml collagen hydrogels showed we could replicate the microenvironment stiffness using these collagen properties. In our findings, using a 7 mg/ml collagen concentration, the average compression modulus was found to be 2.70 ± 0.22 kPa for 5% preload strain. The reported collagen compression moduli for this study are also consistent with our previously reported collagen compression modulus values^26^. In the same study, we also showed that collagen concentration does not alter the diffusivity of solutes, thereby demonstrating that differential response to chemotherapy is not likely due to physical diffusion differences of the collagen concentrations ^26^.

### 3.3 Matrix Stiffness and Hypoxia Modulate Doxorubicin Chemoresistance in Hepatic 2D and 3D Cultures

HCC cell viability in response to doxorubicin treatment under different microenvironmental conditions was measured. The resulting half-maximal inhibitory concentrations (IC_50_) of doxorubicin was quantified for HepG2, C3Asub28, HuH-7, and Hep3B2 cells in 2D monolayers and 3D collagen hydrogels at 4 mg/mL (3D-normal) or 7 mg/mL (3D-cirrhotic) to demonstrate the impact of the tumor structural microenvironment on the hepatic response to this drug. Furthermore, the impact of oxygen concentration was quantified through the environmental regulation of normoxic (21% O_2_) and hypoxic (1% O_2_) conditions for 2D and 3D cultures. The IC_50_ values of doxorubicin after 24-hour treatments are summarized in Figure 3. IC_50_ values for 48-hour treatment durations can be found in Fig S-I and Tables S-II and S-III as minimal changes to IC_50_ values were observed with the increased treatment duration. Overall, we demonstrated that the IC_50_ values of doxorubicin against C3Asub28 cells were the highest compared to other HCC cells, in agreeance with the sub-strain’s increased drug metabolism through CYP3A4.

**Figure 3:**
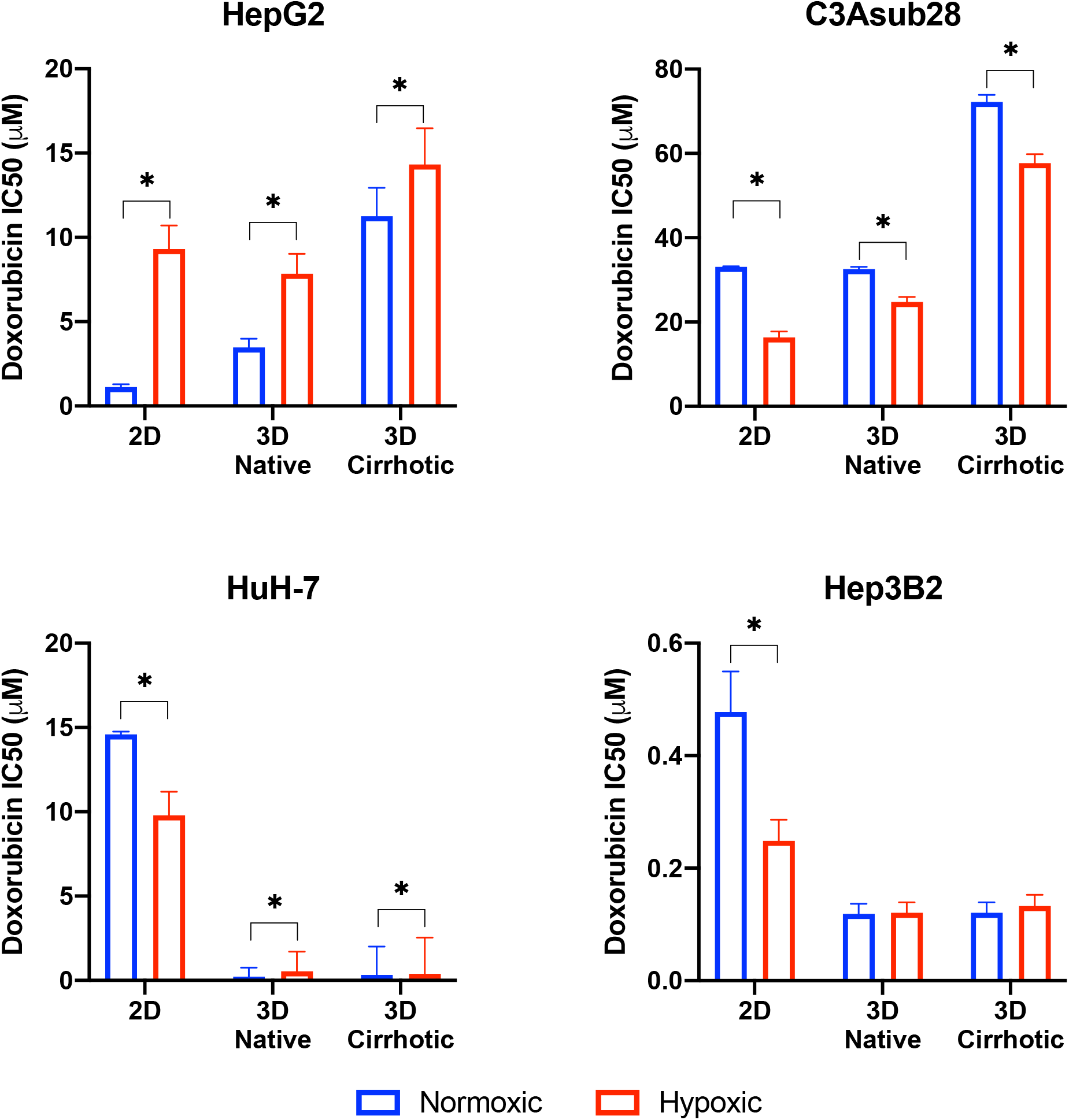
Half-maximal inhibitory concentrations (IC_50_) values of doxorubicin against HCC cells in different microenvironments. HCC cells have stiffness and oxygen-dependent resistance/sensitivity to 24-hour doxorubicin treatment. Cirrhosis and hypoxia altered IC_50_ findings differently among the HCC cell lines. 3D-normal: 4 mg/ml collagen gel, 3D-Cirrhosis: 7 mg/ml collagen gel. Statistical significance was compared to normoxic conditions. *p<0.05.

### 3.4 Comparison of Doxorubicin IC_50_ for 2D vs. 3D Culture under Normoxic Conditions

Studies have demonstrated that 3D matrix stiffness plays an important role in modulation of the cellular response to chemotherapeutics in some cancer cell lines, providing a significantly different response compared to standard 2D culture methods ^27,45^. However, few groups demonstrate the variation of IC_50_ values of doxorubicin between standard 2D-monolayer and 3D culture under varying microenvironmental conditions. In the context of the 3D microenvironment, we were able to determine the influence of normal liver stiffness (4 mg/mL collagen) under normoxic conditions (21% O_2_) compared to 2D-normoxic monolayers as calculated by the fold change in chemoresistance due to the presence of the 3D microenvironment. In general, we demonstrate that IC_50_ values of doxorubicin against HepG2 increased in 3D compared to 2D but did not change for the C3Asub28 cell line. In contrast, 3D culture decreased IC_50_ values of doxorubicin for HuH-7 and Hep3B2 cell lines compared to 2D monolayers.

We observed that IC_50_ of HepG2 cells after a 24-hour doxorubicin treatment was 3.09-fold higher in the 3D normal-normoxic environment (IC_50_ = 1.13 μM) compared to the 2D-normoxic (IC_50_ = 3.48 μM). The IC_50_ of doxorubicin against C3Asub28 cells was consistently the highest, but it was the only cell line that did not lead to a significant change in IC_50_ of doxorubicin in response to the 3D normal-normoxic compared to 2D-normoxic. The efficacy of doxorubicin on HuH-7 cells cultured in 3D normal-normoxic microenvironment was higher than the HepG2 and C3Asub28 phenotypes. Overall, we observed that the IC_50_ of doxorubicin against HuH-7 cells was lower (p<0.05) in 3D normal-normoxic compared to 2D-normoxic by 62.61-fold. Conversely, IC_50_ values of doxorubicin against Hep3B2 cells were the lowest overall and significantly decreased 4.02-fold 3D normal-normoxic compared to 2D-normoxic.

### 3.5 The Influence of Microenvironmental Stiffness on Doxorubicin Chemoresistance under Normoxic Conditions

To isolate the impact that microenvironmental stiffness plays in the regulation of chemoresistance of different HCC cell lines, we analyzed the impact that the shift from normal (4 mg/mL) to cirrhotic (7 mg/mL) collagen concentration has on doxorubicin IC_50_ values under normoxic conditions for a 24-hour treatment duration. Overall, we found that the increase in microenvironmental stiffness, modeled by the higher collagen concentration, increased the IC_50_ values of doxorubicin against the HCC cell lines, HepG2 and C3Asub28, that had a higher basal chemoresistance to doxorubicin. Variations in fold changes for 24-hour treatment were reported in Figure S-II. 48-hour treatment duration resulted in marginal differences in IC_50_ values compared to 24-hour treatment duration (Figure S-III). The IC_50_ of doxorubicin against HepG2 (3.24-fold) and C3Asub28 (2.22-fold) cells cultured in 3D cirrhotic-normoxic conditions increased compared to cells cultured in 3D normal-normoxic conditions. However, for both HuH-7 and Hep3B2.17, we did not see any significant difference in IC_50_ values of doxorubicin against these cells (p>0.05), between 3D normal-normoxic and 3D cirrhotic-normoxic conditions.

### 3.6 Influence of Hypoxia on Doxorubicin Chemoresistance

Oxygen concentration in HCC liver tumors can change due to reduced blood flow, increased cell density, and environmental stiffening ^17,23^. This has been shown to alter both the proliferation rate and chemoresistance of tumor cells. We isolated and quantified the influence of hypoxia on chemoresistance, as measured by IC_50_ values, when cells were cultured in 3D with normal liver stiffness (4 mg/ml) and in 2D monolayers both under hypoxic conditions. Overall, the introduction of hypoxic conditions to 2D monolayers or 3D collagen hydrogels with normal stiffness (4 mg/ml) showed a change in response between the HCC cell lines for 24-hour treatment durations and saw minimal changes for 48-hour treatment durations as shown in Figure S-III. Hypoxia consistently increased IC_50_ values of doxorubicin against HepG2 cells but decreased IC_50_ values of doxorubicin against C3Asub28 IC_50_ compared to normoxic conditions for all stiffnesses. The response of HuH-7 and Hep3B2 cells to doxorubicin under hypoxia was variable depending on if they were cultured in 2D or 3D.

We next isolated the impact that hypoxia plays in the regulation of chemoresistance of different HCC cell lines in 3D microenvironments. We first analyzed the effect that the shift from 4 mg/mL to 7 mg/mL collagen concentration had on doxorubicin IC_50_ values under hypoxic (1% O_2_) conditions and then quantified the impact that the shift from normoxia to hypoxia had within each collagen concentration. Within both normal (4 mg/mL) and cirrhotic (7 mg/mL) liver stiffness, the introduction of hypoxia resulted in a general increase in doxorubicin IC_50_ values, except for C3Asub28 which shows the opposite trend. However, we did not observe a statistical significance of doxorubicin IC_50_ values when Hep3B2 cells cultures in 3D normal-hypoxic or 3D cirrhotic-hypoxic conditions relative to their normoxic counterparts. All of the observed trends in IC_50_ fold changes suggested that HCC cells, depending on the underlying phenotype (denoted by the different HCC cell lines utilized), can display a differential change in IC_50_ values of doxorubicin dependent upon microenvironmental stiffness and oxygen availability.

### 3.7 Matrix Stiffness and Hypoxia Modulate Cell Viability in Response to Sorafenib Treatment in Hepatic 3D Cultures

Sorafenib is not primarily utilized to induce direct cell death, its mechanism is to inhibit kinases responsible for promoting angiogenesis and cell growth, mainly VEGFR, PDGFR, and RAF^15^. As such, even high doses are not sufficient to terminate 50% of the HCC population rendering the calculation of IC_50_ values impossible. We investigated the direct cytotoxic impact of sorafenib to establish baseline values for more clinically relevant combined therapeutic administration of sorafenib and doxorubicin. Clinically relevant standard (11 μM) and high (22 μM) doses of sorafenib reported in previous studies were tested on four different HCC cell lines ^42^. We observed many of the same trends previously observed for doxorubicin chemoresistance (Figure 4). We note that C3Asub28 demonstrated the lowest levels of cell death. We also demonstrate that in C3asub28 cells that an increase in microenvironmental stiffness correlated to a general increase in sorafenib chemoresistance. However, the effects of hypoxia were more varied, and sorafenib chemoresistance differed depending on the cell line, sorafenib dose, and microenvironmental stiffness. In our comparison of sorafenib treated cells in 3D to the untreated 3D controls, we established that cell viability is not significantly impacted at standard, 11 μM, Sorafenib doses for many of the cell lines regardless of matrix stiffness or oxygen concentration at 24-hour treatment durations. However, higher doses of sorafenib, 22 μM, shows a much higher direct cytotoxic effect across all HCC cell lines and microenvironmental conditions tested. The influence of hypoxia on sorafenib chemoresistance has a markedly different dynamic than what we have previously seen with doxorubicin.

**Figure 4:**
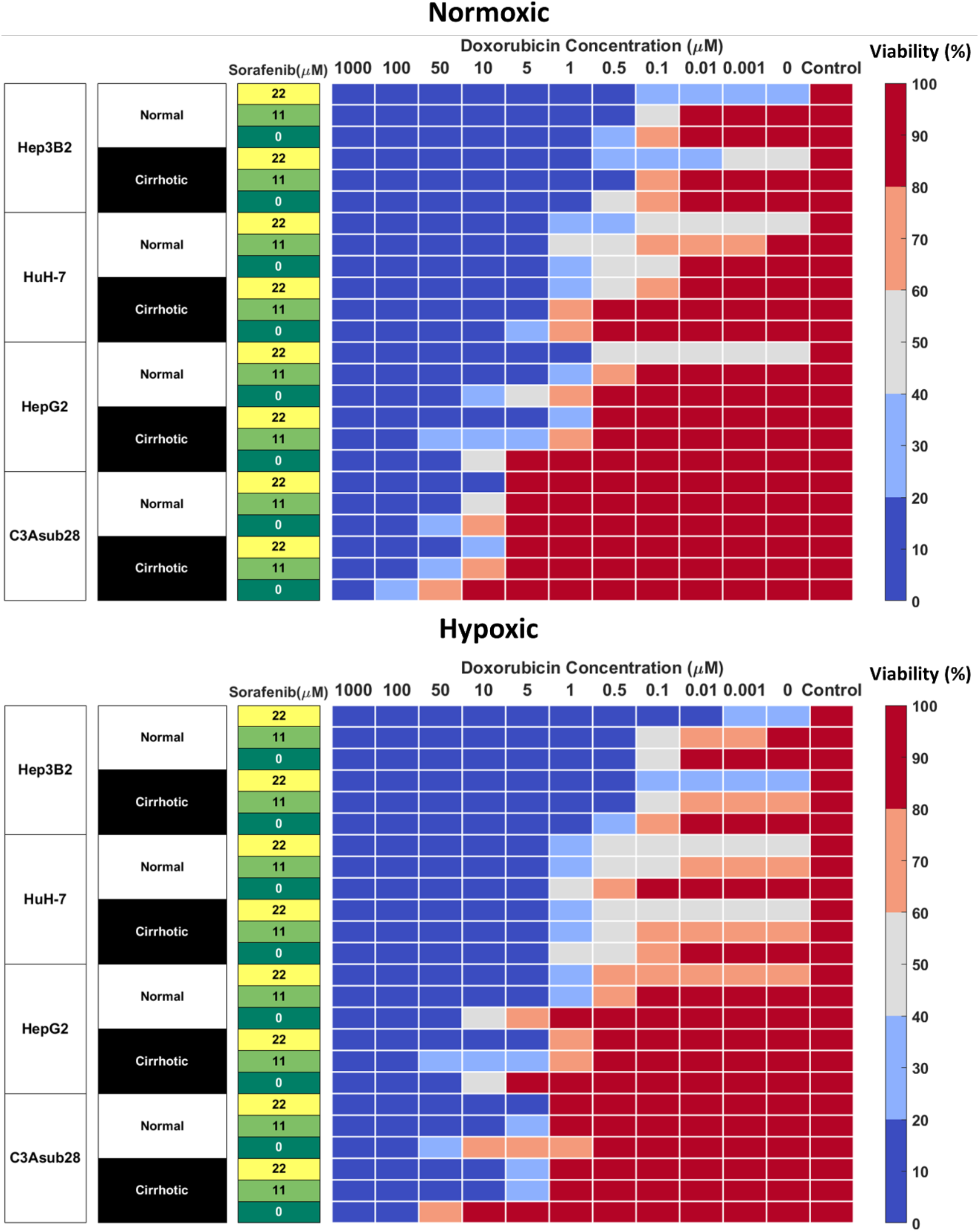
Combined doxorubicin and sorafenib treatment efficacy of HCC cell lines under the influence of hypoxia and cirrhosis. HCC cell types show sensitivity to combined sorafenib and doxorubicin treatment for 24 hours. Cells cultured in 3D normal and cirrhotic hydrogels in hypoxic and normoxic conditions. Cell viability was analyzed and plotted as a percentage of the untreated control. A combination of doxorubicin with sorafenib improves treatment efficacy. The influence of cirrhosis decreased HCC cell death.

### 3.8 Sorafenib Improves Minimum Required Dose of Doxorubicin for Acute Toxicity

Combined administration of doxorubicin and sorafenib in HCC patients has been commonly used due to their potential synergistic effect ^41,46^. Doxorubicin is used to inhibit tumor proliferation^47^, while sorafenib is used to inhibit angiogenesis and tumor cell proliferation in the TME ^15^. Cells cultured in 3D normal and cirrhotic hydrogels were investigated for the impact of doxorubicin-sorafenib combination therapy to determine how the introduction of sorafenib influenced the resulting cell viability. Cell viability in normal and cirrhotic tissue with a 24-hour treatment duration of both drugs and the influence of hypoxia are presented in Figure 4 and changes in doxorubicin IC_50_ values due to the presence of sorafenib discussed in this section. Treatment efficacy is reported as fold viability change compared to untreated controls reported in Figure S-IV. For this analysis, statistical comparison (p<0.05) of viability under tested doses and untreated samples were compared, and minimum dose to induce cytotoxicity was reported. We found minimal differences between 24- and 48-hour treatments with these drugs and present 48-hour treatment results in Figure S-V.

HepG2 cells treated with sorafenib combination therapy demonstrated a general decrease in doxorubicin IC_50_. Under 3D normal-normoxic conditions, the doxorubicin IC_50_ values decreased from 3.48 μM to 0.68 μM (p=0.01) and 0.02 μM (p=0.005) for combination therapy with standard and high doses of sorafenib, respectively. Similarly, 3D cirrhotic-normoxic conditions with sorafenib combination therapy reduced the doxorubicin IC_50_ from 11.25 μM to 2.42 μM (p=0.04) and 0.92 μM (p=0.02) for standard and high doses of sorafenib, respectively. Under hypoxic conditions, we found that the trend held. HepG2 cells in the normal-hypoxic conditions in combination therapy showed decreased doxorubicin IC_50_ values from 7.85 μM to 0.71 μM (p=0.03) and 0.31 μM (p=0.01) for combination therapy with standard and high doses of sorafenib, respectively. Under cirrhotic-hypoxic conditions, combination therapy with sorafenib reduced doxorubicin IC_50_ values from 14.33 μM to 2.9 μM (p=0.05) and 2.85 μM (p=0.05) for combination therapy with standard and high doses of sorafenib, respectively. Although the HepG2 cells showed a higher chemoresistance to doxorubicin in hypoxic and cirrhotic culture conditions relative to normoxic and normal, the combination with sorafenib significantly diminished doxorubicin IC_50_ values for all conditions tested.

The C3Asub28 cell line demonstrated the highest levels of chemoresistance out of all of the selected HCC cell lines, however consistent with other cell lines, combination therapy with sorafenib considerably reduced doxorubicin IC_50_ values for all conditions. Under 3D normal-normoxic conditions, the doxorubicin IC_50_ values decreased from 32.58 μM to 9.81 μM (p=0.04) and 8.36 μM (p=0.006) for combination therapy with standard and high doses of sorafenib, respectively. Similarly, 3D cirrhotic-normoxic conditions with sorafenib combination therapy reduced the doxorubicin IC_50_ from 72.23 μM to 22.25 μM (p=0.05) and 9.46 μM (p=0.02) for standard and high doses of sorafenib, respectively. Under the normal-hypoxic condition, cells in combination therapy showed decreased doxorubicin IC_50_ values from 24.76 μM to 3.99 μM (p=0.05) and 3.66 μM (p=0.04) for standard and high doses of sorafenib, respectively. Under cirrhotic-hypoxic conditions, combination therapy with sorafenib reduced doxorubicin IC_50_ values from 57.68 μM to 4.48 μM (p=0.05) and 3.78 μM (p=0.05) for combination therapy with standard and high doses of sorafenib, respectively.

HuH-7 cells under the normal-normoxic condition showed a decrease in doxorubicin IC_50_ values from 0.43 μM to 0.33 μM and 0.01 μM (p=0.009) for combination therapy with standard and high doses of sorafenib, respectively. Similarly, 3D cirrhotic-normoxic conditions with sorafenib combination therapy reduced the doxorubicin IC_50_ from 2.24 μM to 1.61 μM and 0.31 μM (p=0.03) for standard and high doses of sorafenib, respectively. Under hypoxic conditions, we found that HuH-7 cells in the normal-hypoxic conditions showed decreased doxorubicin IC_50_ values from 0.54 μM to 0.24 μM (p=0.05) and 0.02 μM (p=0.005) for combination therapy with standard and high doses of sorafenib, respectively. Under cirrhotic-hypoxic conditions, combination therapy with sorafenib reduced doxorubicin IC_50_ values from 0.4 μM to 0.25 μM (p=0.05) and 0.01 μM (p=0.008) for combination therapy with standard and high doses of sorafenib, respectively.

For Hep3B2 cells, high doses of sorafenib (22 μM) were sufficient to reduce cell viability below 50% without the addition of doxorubicin in all conditions tested. However, combination therapy with standard doses of sorafenib (11 μM) reduced doxorubicin IC_50_ values in all conditions tested. Under 3D normal-normoxic conditions, the doxorubicin IC_50_ values decreased from 0.12 μM to 0.1 μM for combination therapy with standard doses of sorafenib. Similarly, 3D cirrhotic-normoxic conditions with sorafenib combination therapy reduced the doxorubicin IC_50_ from 0.3 μM to 0.14 μM (p=) standard doses of sorafenib. Under the normal-hypoxic conditions, combination therapy showed decreased doxorubicin IC_50_ values from 0.12 μM to 0.04 μM (p=0.01) for combination therapy with standard doses of sorafenib. Under cirrhotic-hypoxic conditions, combination therapy with sorafenib reduced doxorubicin IC_50_ values from 0.13 μM to 0.04 μM (p=0.02) for standard doses of sorafenib.

### 3.9 Cirrhosis and Hypoxia Regulated HCC Metabolic Activity

The CYP3A4 enzyme is one of the major mechanisms of drug metabolism for cancer therapeutics, including sorafenib and doxorubicin, in the liver. These enzymes can metabolize drugs before they have the chance to cause their intended direct cytotoxic effects on the cells.^14^. CYP3A4 metabolic activity of the HCC cell lines was measured in 2D monolayers and in 3D collagen I hydrogels in response to different stiffness and hypoxic conditions. Regulation of CYP3A4 by cirrhosis and hypoxia is presented in Figure 5. In agreement with the previously published literature, the C3Asub28 cell line expressed much higher CYP3A4 expression than other HCC cell lines^38^. In our study, C3Asub28 CYP3A4 expression is 7.17 ± 2.73 fold higher than the HepG2 cell line, which is within the range of previously published work (6.1 ± 0.2 fold) ^38^. The introduction of hypoxia significantly downregulated CYP3A4 expression compared to normoxia in all microenvironments (p=0.03). CYP3A4 expression of C3Asub28 cells did not change with culture in 3D normal-normoxic and 3D normal-hypoxic conditions than similar 2D conditions (p=0.78).

**Figure 5:**
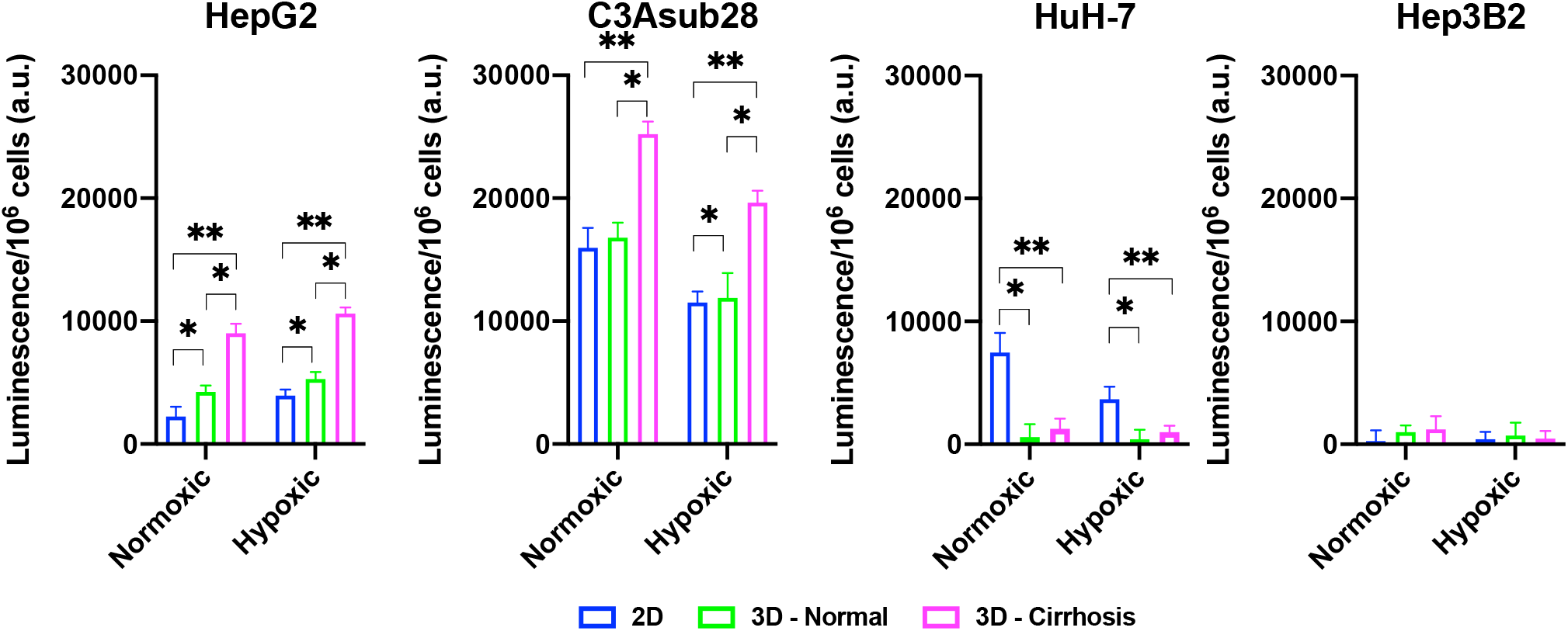
Regulation of CYP3A4 is responsible for drug metabolization by HCC cell lines in response to hypoxic and cirrhotic conditions. Presence of cirrhosis and hypoxia altered CYP3A4 activity differently between HCC cell lines. Data represented as mean ± standard deviation. * denotes significance for cirrhotic and normal 3D samples compared to the 2D monolayer. * p < 0.05, ** p < 0.01.

On the other hand, 3D cirrhotic-normoxic and cirrhotic-hypoxic culture upregulated CYP3A4 expression by 1.50 (p=0.007) and 1.65 (p=0.01) fold compared to 3D normal-normoxic and normal-hypoxic conditions, respectively. HepG2 cell lines expressed significantly lower (p=0.02) CYP3A4 expression compared to the C3Asub28 cell line. However, unlike the C3Asub28 cell line, hypoxia upregulated CYP3A4 expression in HepG2 cell lines compared to normoxia for both 2D and 3D microenvironments (p=0.04). 3D normal-normoxic and normal-hypoxic culture upregulated CYP3A4 expression compared to 2D normoxic and 2D hypoxic culture by 1.90 (p=0.03) and 1.35 (p=0.003) fold, respectively. Similarly, 3D cirrhotic-normoxic and cirrhotic-hypoxic culture upregulated CYP3A4 expression compared to 3D normal-normoxic and normal-hypoxic stiffness by 2.13 (p=0.004) and 2.00 (p=0.0002) fold, respectively. HuH-7 cells expressed relatively lower values of CYP3A4, and none was detected in 3D culture conditions. Furthermore, the Hep3B2 cell line did not express any CYP3A4 expression in all culture conditions. These results show that HCC cells express different basal levels of CYP3A4 and that this expression is can be directly regulated by TME oxygen concentration and stiffness.

Lastly, Pearson correlation between measured CYP3A4 expression and doxorubicin IC_50_ findings with or without the influence of sorafenib was analyzed and presented in Figure 6. Accordingly, we observed a strong correlation between CYP3A4 metabolite and IC_50_ findings for HepG2, C3Asub28, and Hep3B2 cell lines. However, this correlation was not observed for HuH-7 cell line. In this study, we found only HepG2 and C3Asub28 cell lines were regularly expressing CYP3A4 expression in 3D culture conditions unlike to HuH-7 and Hep3B2. For that reason, only HepG2 and C3Asub28 cell lines’ CYP3A4 expression regulation should be taken into account in alternation of IC_50_ values. For that reason, we should conclude that HepG2 and C3Asub28 cell lines are more appropriate for the investigation of CYP3A4 expression regulation.

**Figure 6:**
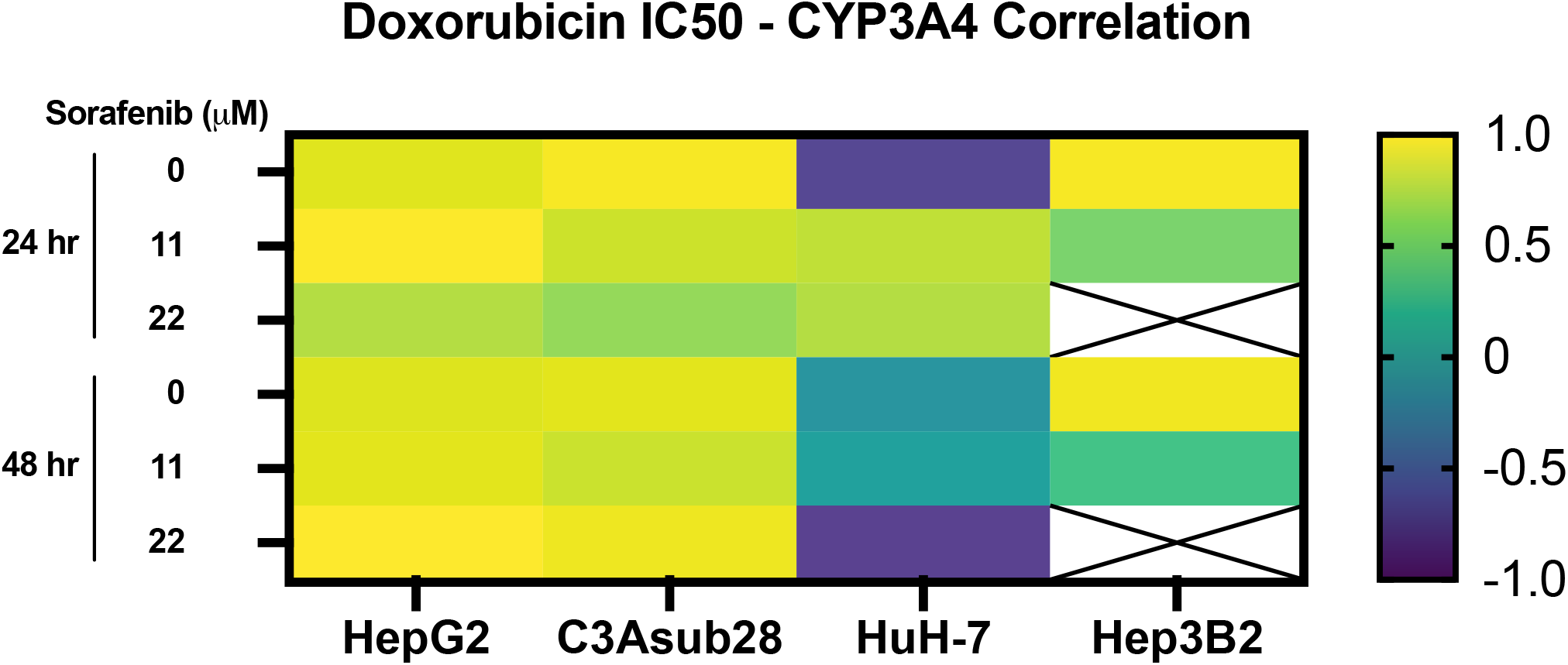
Pearson correlation between CYP3A4 expression and doxorubicin IC_50_ findings with and without the influence of sorafenib treatment in 3D culture conditions. Strong correlation was observed in CYP3A4 expressing HepG2 and C3Asub28 cells unlike to HuH-7 and Hep3B2. Array cells with cross represent IC_50_ values were not detected within tested doses.

## 4 Discussion

In this study, we establish that chemoresistance can be regulated by hypoxic and cirrhotic conditions in the TME through direct modulation of CYP3A4 expression. This regulation can differentially alter the efficacy of chemotherapeutic drugs in HCC cell lines, which potentially has clinical translation to patient-specific HCC treatments. We examined the direct impact of TME stiffness and oxygen concentration, variables commonly associated with HCC tumors on cellular response to chemotherapeutics. This impact was measured by determining the difference in cell viability in response to chemotherapeutic treatment and regulation of the expression of the primary drug-metabolizing enzyme CYP3A4. Using a collagen-based hydrogel system, we observed that 3D culture alone significantly modulates resistance to doxorubicin and sorafenib in HepG2, C3Asub28, and HuH-7 cells but not for Hep3B2 cells. This stiffness-dependent resistance was not observed in similar Hep3B2 cultures, which warrants further investigation into potential phenotypic and genotypic differences in this specific cell line that might elucidate this response. This differential regulation of chemoresistance in different cell lines of the same cancer type is not unique to HCC. This phenomenon has also been observed in other cancers: The chemoresistance indicator, ethoxyresorufin, was upregulated in 3D culture conditions compared to 2D culture in the C3A cell line. This is in agreement with HepG2 and C3Asub28 doxorubicin treatment findings ^48^. Equivalently, the IC_50_ of C3A cells after treatment with paracetamol, trovafloxacin, and fialuridine were found to be higher in 2D culture conditions compared to 3D culture conditions. This result corresponds with our HuH-7 and Hep3B2 doxorubicin chemoresistance findings ^24^. Overall, these studies suggest that drug efficacy in 2D vs. 3D conditions depends on cell phenotype and drug type. This coincides with our findings in which HepG2 and C3Asub28 cell lines had greater chemoresistance in 3D culture compared to 2D but showed the converse trend for HuH-7 and Hep3B2 cells.

Cirrhosis and desmoplastic stiffening in the TME has been shown to be potential factors regulating drug chemoresistance ^49^. In our study, HepG2 chemoresistance to sorafenib and doxorubicin increased in response to cirrhotic stiffness relative to their culture in a matrix of normal stiffness. The C3Asub28 cell line demonstrated a higher chemoresistance to doxorubicin in response to cirrhotic conditions than other tested HCC cells. Similar to the HepG2 cell line, a rise in stiffness also increased the chemoresistance to doxorubicin of the C3Asub28 cell line. However, sorafenib alone did not alter the cell viability of the C3Asub28 cell line for the considered doses potentially attributed to the cell line’s high baseline CYP3A4 metabolic activity. The resistance of HuH-7 and Hep3B2 cells to doxorubicin was not altered in response to cirrhosis, whereas CYP3A4 expression of these cell lines did not change in response to cirrhosis. However, the same cell lines were shown to have higher chemoresistance to sorafenib in response to cirrhosis. Although CYP3A4 carries out the majority of metabolic activity in hepatocytes, other minor cytochromes, such as CYP1A2, 2A6, 1A2, and 2C9, exist and may also alter the drug chemoresistance to an as yet unknown extent. The additional CYP expression present in HCC cells may be responsible for the differential effect of cirrhosis on chemoresistance differences between sorafenib and doxorubicin ^50^. Furthermore, studies in literature showed that a rise in stiffness does not always increase the chemoresistance of cancer cells ^49^. The increase in TME stiffness may improve IC_50_ of MDA-MB-231 triple-negative breast cancer cells to doxorubicin ^27^. However, the same study showed that MCF-7 HER2+ breast cancer cells did not show a stiffness-dependent resistance to doxorubicin. This study hypothesized the increase of stiffness altered chemoresistance differently because MCF-7 remained in an epithelial phenotype, but MDA-MB-231 had a mesenchymal phenotype. Similarly, stiffness induces chemoresistance of BxPC-3 and Suit2-007 pancreatic cancer cells to paclitaxel, but not to gemcitabine ^51^. These studies hypothesized the differential effect of chemoresistance to different drugs between cell lines could be related to phenotypical differentiation from epithelial to mesenchymal phenotype. In addition, HepG2, HuH-7, and Hep3B2 cell lines have different phenotypic profiles and differentiation levels, potentially explaining differences in their chemoresistance and metabolic activity in response to cirrhosis ^52^.

Hypoxia is known to be one of the regulating factors of chemoresistance ^53^. In our study, HepG2 chemoresistance to doxorubicin and sorafenib increased in response to hypoxia compared to the normoxic condition as measured both by increased IC_50_ values and CYP3A4 expression. However, for C3Asub28 and HuH-7, chemoresistance decreased in response to hypoxia compared to normoxia. Previous studies have also demonstrated the differential effect of hypoxia on drug efficacy, depending on the cell line and phenotype. The presence of hypoxia has been shown to upregulate hypoxia-induced factor (HIF1-α), but this alters the CYP isoforms differently in various medulloblastoma cell lines ^54^. The molecular pathway still could not be explained in this study, but it has been hypothesized that nuclear receptors, namely PPARα, PPARγ, or ER-α, as well as the constitutive androstane and pregnane X receptors, have found to be altered differently under hypoxia ^54^. In addition to this, chemoresistance does not always increase in response to hypoxia ^53^. The same study also showed the regulation of chemoresistance under hypoxia is not universal between ovarian, renal, breast, lung, and lymphoma cancer cell lines and varies for different drugs. This supports our data showing the differential effects of hypoxia on doxorubicin IC_50_ values of the tested HCC cell lines ^53^. Also, hypoxia increases HepG2 chemoresistance to doxorubicin, in confirmation with our study, but not to rapamycin ^23^. In parallel to this, a significant decrease in apoptotic cells induced by cisplatin was reported under hypoxic conditions for HepG2 and MHCC97L cell lines, which is in agreement with what we observed with increased chemoresistance of HepG2 cells under hypoxic conditions ^55^. It has also been showed that hypoxia downregulates drug-metabolizing enzymes and subsequently the chemoresistance of the HepaRG hepatoma cell line, which agrees with our findings of C3Asub28 CYP3A4 modulation under hypoxia ^56^.

Consequently, the differential role of hypoxia on molecular chemoresistance expressions and drug efficacy has been reported. It has been shown that HIF1-α is upregulated due to a lack of oxygen in TME. This may or may not induce drug transporters such as MDR1 and targets of delivered drugs (topoisomerase II) in each cell line ^20^. Additionally, possible nuclear receptors have been proposed to regulate CYP3A4 in response to hypoxia through the expression of HIF1-α and p53 expression^57^. Alternation of molecular drug transport mechanisms could be the reason why we observed variable chemoresistance between different HCC cell lines under hypoxia.

Our study showed CYP3A4 expression is regulated by microenvironment stiffness and hypoxia for HepG2, C3Asub28, and HuH-7 cell lines providing a potential mechanism connecting the TME to the chemotherapeutic response. The regulation of CYP3A4 resulted in a significant impact on the efficacy of doxorubicin and sorafenib, whose trends in the regulation mirror the observed changes in cell viability in response to the drugs. Doxorubicin IC_50_ was higher, and sorafenib terminated less HCC population for HepG2 and C3Asub28 cells in cirrhotic, 7 mg/ml, microenvironments in general compared to healthy, 4 mg/mL, stiffness reflecting the CYP3A4 expression in those microenvironments. However, we did not see any significant IC_50_ change in response to doxorubicin for the HuH-7 cell line when cultured in normal and cirrhotic 3D microenvironments. In addition, CYP3A4 expression of Hep3B2 did not change when cultured in normal and cirrhotic 3D microenvironments, which is parallel with IC_50_ findings.

Moreover, in general, we demonstrate that hypoxia increases doxorubicin IC_50_ against HepG2 cells but not for C3Asub28 cells. The change of doxorubicin IC_50_ was parallel to the regulation of CYP3A4 expression. Hypoxia upregulated CYP3A4 expression of HepG2 cells but downregulated CYP3A4 expression of C3Asub28 cells. This work further confirms the upregulated metabolic expression of CYP3A4 decreases doxorubicin efficacy in confirmation with previous studies across multiple cancer types, including liver^58^, colorectal^29^, breast^59^, and prostate^60^. Inhibition of CYP3A4 activity of human primary hepatocytes has shown to suppress human pregnane X receptor (hPXR) agonist-induced chemoresistance ^61^. Overexpression of CYP3A4 in tumor tissues and chemoresistance to therapeutics has been shown in clinical practices ^62^. Similarly, there is evidence that therapeutic efficacy of drugs have been diminished through CYP3A4 enzyme expressed by hepatocellular carcinoma cells^63^. These studies agree with our findings on the regulation of chemoresistance based on CYP3A4 activity. However, our work presents the significant finding that the regulation of CYP3A4 expression can be directly tied to the tumor microenvironment. We demonstrate that CYP3A4 activity can be regulated by oxygen concentration and TME stiffness, subsequently altering the metabolism of the chemotherapeutic drugs in HCC cell lines. Possible nuclear receptor pathways regulating CYP3A4 in response to hypoxia through HIF1-α, p53, PPARα, VDR, FXR, and LXR have been proposed and hypothesized that hypoxia could affect CYP3A4 at different degrees ^57^. Furthermore, TME stiffness has been hypothesized to alter CYP3A4 differentially through yes-associated protein (YAP) pathway ^64^. This likely has a direct clinical translation to *in vivo* HCC regulation enforced by the clinical observations of highly variable patient to patient HCC CYP3A4 expression ^65^. The majority of the current HCC treatments result in poor treatment outcomes, as such, the consideration of tumor microenvironment properties (such as stiffness variation due to the changing fibrosis scores of patients, presence of different levels of hypoxia as a result of this desmoplastic stiffening in the TME), and CYP expression levels could potentially bring benefits to outcomes of HCC treatment, and provide a basis for personalized HCC treatment ^62,66^.

Overall, this work demonstrates TME stiffness and oxygen concentration modulates CYP3A4 expression of HCC cells and, consequently, their chemoresistance to doxorubicin and sorafenib treatment. We determined the existence of a stiffness-dependent resistance to doxorubicin and sorafenib, depending on the differential genetics of the HCC, such as phenotypical changes from epithelial to mesenchymal ^27^. HepG2 and C3Asub28 cells showed a higher chemoresistance to doxorubicin and sorafenib under cirrhotic conditions. Conversely, we did not observe a change in chemoresistance for HuH-7 and Hep3B2 cell lines to doxorubicin in response to cirrhotic conditions, which may be due to underlying genotypic differences including differentiation levels, which alter metabolism pathways through glucose, glutamine, and glutamate ^52^. Hypoxia demonstrated a much different impact on the HCC cells, upregulating the chemoresistance of HepG2 and HuH-7, but downregulating chemoresistance of the C3Asub28 cell line. We did not observe a significant variation of chemoresistance in the Hep3B2 cell line in 3D culture. Drug metabolism, measured by CYP3A4 expression, mirrored effective chemoresistance measured by IC_50_ values in cell vitality assays. We saw an increase in CYP3A4 expression in the 3D culture of C3Asub28 and HepG2 cell lines compared to 2D, but this expression decreased for the HuH-7 cell line for both hypoxic and normoxic conditions. At a minimum, the presence of a 3D culture system significantly modulates the response of HCC to chemotherapy over the standard 2D methods. Previously it has been shown 3D culture alters the integrin ligands (such as AKT and RAF), which are the targeting for doxorubicin and sorafenib, differently among different cell lines compared to 2D culture ^49^.

Further, the stiffness of the microenvironment, seen in HCC patients with cirrhosis, modulates drug resistance and should be taken into consideration when determining treatment options and doses. While CYP3A4 maintains the majority of the drug metabolism in the liver, other enzymes such as CYP2A6, 1A2, and 2C9 has shown to have a minor contribution to drug metabolism ^62^, and provide further insight in IC_50_ of Hep3B2 variation we observed between 2D and 3D microenvironments but not captured a difference in CYP3A4 expression. Further expansion of this work is needed to investigate the varied response between existing HCC lines or patient-specific primary tumor cells that might provide insight into the known differential effectiveness of standard HCC treatments *in vivo* ^66^. Potentially, other drugs used to treat HCC in clinical practice such as lenvatinib could be tested in this system to observed its potential effect on HCC treatment ^67^.

HCC is diagnosed based on imaging and laboratory criteria, making it the only cancer that is diagnosed without biopsy. This practice has come under increasing scrutiny for many reasons, especially given the growing appreciation of the value of precision medicine. Moreover, while some progress has been made, HCC still responds very poorly to drugs in general and notably to immunotherapies. The results from the present study underscore the importance of restoring the practice of obtaining biopsy specimens to obtain the necessary information. Despite much effort to identify characteristic biomarkers (both serum and newer methods of elastography), important gaps remain. An understanding of the degree of fibrosis, baseline expression of CYP3A4, and the immune landscape will become essential in developing drug treatment plans and clinical trials.

## 5 Acknowledgment

The authors would like to thank Dr. Wei Li (Department of Mechanical Engineering, the University of Texas at Austin) and Dr. Christopher Sullivan (Department of Molecular Sciences) for generously gifting C3Asub28 cells and HuH-7, respectively. This work was completed with support from the Veterans Health Administration and with resources and the use of facilities at the Central Texas Veterans Health Care System, Temple, Texas. The contents do not represent the views of the U.S. Department of Veterans Affairs or the United States Government. T.E.Y. is a CPRIT scholar in cancer research.

## 7 Author contributions

Conceptualization: A.O. and M.N.R. conceived of the idea for the study. Supervision: E.N.K.C, T.E.Y. and M.N.R. supervised the project. E.N.K.C., S.D., M.M., T.E.Y. and M.N.R provided feedback and assistance with manuscript preparation. Investigation: A.O., D.L.S., and M.N.R were responsible for performing the studies and analyzing the experimental data. Writing: A.O. wrote the initial draft of the paper. All authors discussed the results and revised the manuscript.

